# Inferring phenotypic plasticity and local adaptation to climate across tree species ranges using forest inventory data

**DOI:** 10.1101/527390

**Authors:** Thibaut Fréjaville, Bruno Fady, Antoine Kremer, Alexis Ducousso, Marta Benito Garzón

## Abstract

**Aim:** To test whether adaptive and plastic trait responses to climate across species distribution ranges can be untangled using field observations, under the rationale that, in natural forest tree populations, long-term climate shapes local adaptation while recent climate change drives phenotypic plasticity.

**Location:** Europe.

**Time period:** 1901-2014.

**Taxa:** Silver fir (*Abies alba* Mill.) and sessile oak (*Quercus petraea* (Matt.) Liebl.).

**Methods:** We estimated the variation of individual tree height as a function of long-term and short-term climates to tease apart local adaptation, plasticity and their interaction, using mixed-effect models calibrated with National Forest Inventory data (*in-situ* models). To validate our approach, we tested the ability of *in-situ* models to predict independently tree height observations in common gardens where local adaptation to climate of populations and their plasticity can be measured and separated. *In-situ* model predictions of tree height variation among provenances (populations of different geographical origin) and among planting sites were compared to observations in common gardens and to predictions from a similar model calibrated using common garden data (*ex-situ* model).

**Results:** In *Q. petraea*, we found high correlations between *in-situ* and *ex-situ* model predictions of provenance and plasticity effects and their interaction on tree height (*r* > 0.80). We showed that the *in-situ* models significantly predicted tree height variation among provenances and sites for *Abies alba* and *Quercus petraea*. Spatial predictions of phenotypic plasticity across species distribution ranges indicate decreasing tree height in populations of warmer climates in response to recent anthropogenic climate warming.

**Main conclusions:** Our modelling approach using National Forest Inventory observations provides a new perspective for understanding patterns of intraspecific trait variation across species ranges. Its application is particularly interesting for species for which common garden experiments do not exist or do not cover the entire climatic range of the species.

## INTRODUCTION

Understanding the causes of phenotypic variation across species distribution ranges is important because phenotypic traits are fundamental drivers of community assembly, ecosystem functioning, and population response to climate change (Diaz *et al.*, 2004; Shipley *et al.*, 2006; Alberto *et al.*, 2013; Kunstler *et al.*, 2016). The two non-exclusive causes of phenotypic variation within species are the phenotypic plasticity (i.e., the capacity of one genotype to render different phenotypes under different environments, Valladares *et al*., 2006) and local adaptation (i.e., the fact that individuals have a better fitness in their local environment than individuals from other populations, Kawecki & Ebert, 2004). The spatial distribution of the amount of phenotypic variation that can be attributed to phenotypic plasticity or to local adaptation may change the response of organisms to climate change as predicted by theoretical approaches (Chevin *et al.*, 2010; Valladares *et al.*, 2014). Yet, attributing the cause of phenotypic trait variation across species ranges to either plastic or genetic components, or the interaction between the two, remains a challenge without using costly, long-term common garden experiments.

Long-term spatial divergence under different environmental conditions is known to promote phenotypic differentiation of populations as a result of local adaptation (Mimura & Aitken, 2010; Savolainen *et al.*, 2013; Yeaman *et al.*, 2016), which affects their current response to a particular climate (Rehfeldt *et al.*, 2002; Savolainen *et al.*, 2007; Valladares *et al.*, 2014). However, significantly less is known about the distribution of phenotypic plasticity and its importance for populations for coping with rapid climate change across the species range, especially in long-lived sessile organisms such as forest trees (Nicotra *et al.*,2010; Benito Garzón *et al.*, 2011; Valladares *et al.*, 2014; Duputié *et al.*, 2015).

Patterns of phenotypic plasticity and local adaptation of populations have long been assessed using common garden or reciprocal transplant experiments, in which genotypes of known climatic origin (i.e., provenances) are growing in experimental plantations where short-term environmental conditions are controlled. In common gardens (also named ‘provenance tests’ or ‘genetic trials’), local adaptation has mostly been inferred from trait differences among provenances that are related to the long-term climate of origin of the provenance, while plasticity is quantified by trait variation with the short-term climatic conditions at the planting sites (see Matyas, 1994; Wang *et al.*, 2006; Leites *et al.*, 2012 for forest trees). Common gardens have been established for a few economically important tree species for which only a restricted range of populations and ontogenic stages have been studied, which makes the understanding of phenotypic variation across species ranges limited (Fady *et al.*, 2016).

On the other hand, causes of phenotypic variation are confounded in natural conditions, in addition to the effects of ontogeny and competition. National Forest Inventories (NFI) provide extensive data of phenotypic variation of forest trees in natural conditions, and hence, they have been widely used to test different ecological questions such as the effects of functional traits on competition, forest productivity and response to climate change (Kunstler *et al.*, 2016; Ratcliffe *et al.*, 2016; Ruiz-Benito *et al.*, 2017), but to date, the causes of phenotypic variation in NFI remain unexplored.

Here we show that the components of intraspecific trait variation: local adaptation, plasticity and their interaction, can be statistically estimated using the field data recorded in French NFI for two ecologically and economically important forest tree species usually managed using natural regeneration, *Abies alba* Mill. and *Quercus petraea* (Matt.) Liebl. We use tree height, an important adaptive and fitness-related trait (Savolainen *et al.*, 2007; Díaz *et al.*, 2016), to independently test our approach on field observations (NFI) and we validate our findings using common garden data. Our approach expands the space-for-time substitution analysis developed in common gardens (Matyas, 1994; Rehfeldt *et al.*, 2002; Leites *et al.*, 2012) to field observations of phenotypic trait variation, with the rationale that trees inventoried in the field have a local origin (i.e. seed sources originated within the bioclimatic region inhabited by the trees). To separate the sources of phenotypic variation in nature, we examined climatic variations that occur at two temporal and spatial scales: first, regional patterns in long-term climate (LTC) that have promoted trait variation among provenances as a result of local adaptation (Savolainen *et al.*, 2007; Mimura & Aitken, 2010; Kremer *et al.*, 2012); and, second, short-term climate (STC) that shapes plastic responses of individual trees to recent climate change (Nicotra *et al.*, 2010; Valladares *et al.*, 2014). Our approach opens new perspectives for the understanding of phenotypic variation patterns across species distribution ranges using large field observation datasets such as forest inventories.

## METHODS

We analysed tree height (m), a fitness-related phenotypic trait, measured both in NFI and common gardens. We selected two major European forest species with contrasted life history traits and ecological requirements: *Abies alba* Mill. (a montane evergreen needle-leaved gymnosperm) and *Quercus petraea* (Matt.) Liebl. (a temperate deciduous broadleaved angiosperm). In the NFI, these two species are traditionally managed using natural regeneration, thus adult trees are assumed to derive from the local gene pool.

We calibrated two independent mixed-effect models of individual tree height using NFI (*in-situ* model) and common garden data (*ex-situ* model), respectively. To validate our models we used two different methods (Fig. 1). The first one is a validation using common garden data: it directly compares the results of the *in-situ* model with independent tree height measurements standardized by common garden and by provenance to respectively separate the effects of the provenance (local adaptation) and plasticity. The second one is a validation using *ex-situ* model predictions: it compares the predictions of *in situ* and *ex-situ* models regarding the relative contribution to the model of the climate of the planting site (plastic effect) and that of the climate of the origin of the provenances (local adaptation), and the interaction between both. All analyses and computations were carried out in the R software environment (R Core Team, 2013).

**Figure 1.**
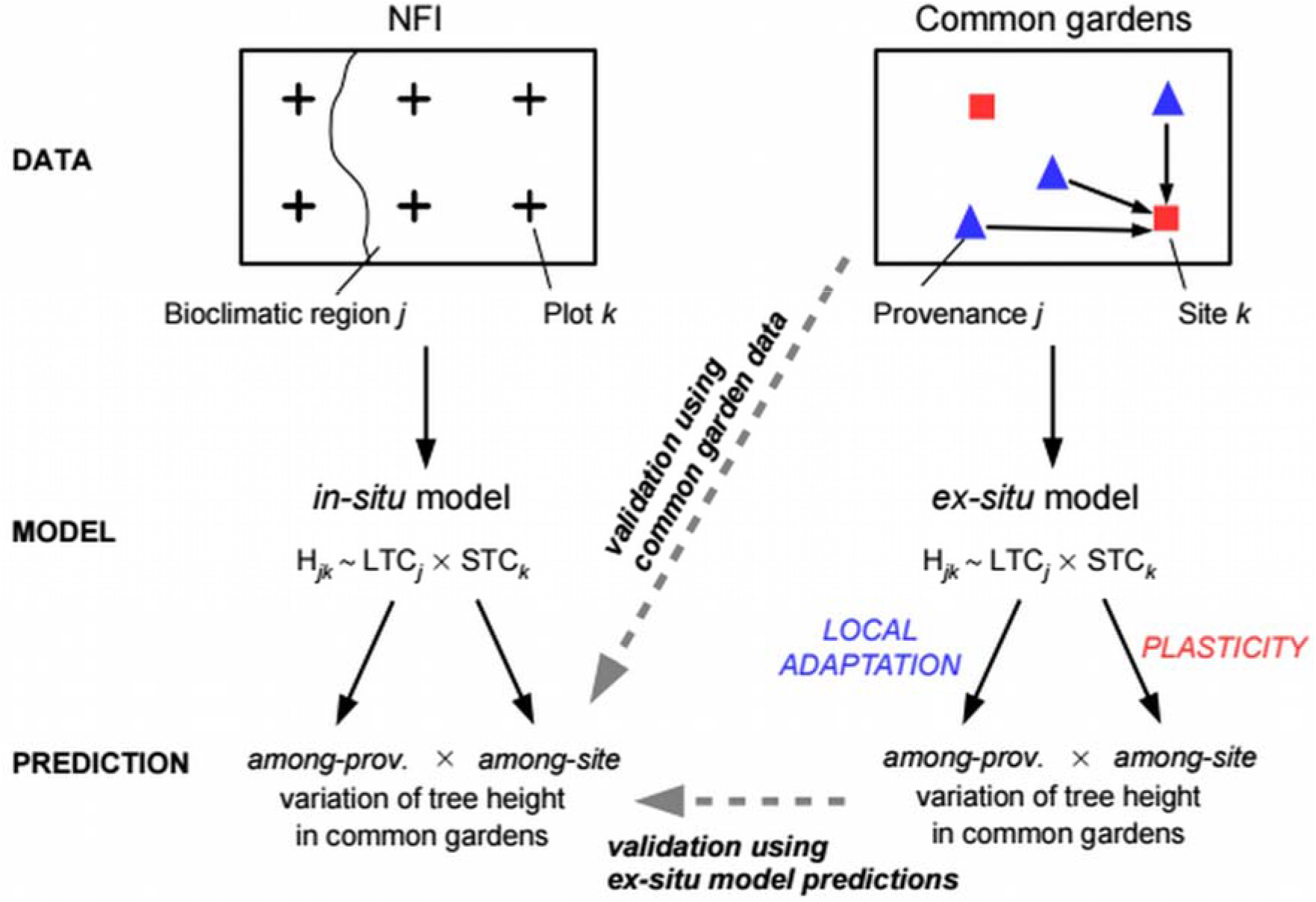
Workflow of the modelling approach and validation methods. We used individual tree height data from National Forest Inventories (NFI) to calibrate a mixed-effect model (‘*in-situ* model’) as a function of long-term climate (LTC) of the bioclimatic region and short-term climate (STC) of the forest plot to disentangle local adaptation, plasticity and their interaction on intraspecific trait variation. To validate our approach, we compared *in-situ* model predictions with independent observations of tree height variation in common gardens where trait differences among populations of different geographical origin (i.e., the provenance) and their plasticity can be separated. *In-situ* model predictions of tree height variation among provenances and among planting sites were compared to observations in common gardens (*validation using common garden data*) and to predictions from a parallel model calibrated using common garden data *(validation using ex-situ model predictions). Validation using ex-situ model predictions* needs common garden data covering large climatic gradients (as is the case of *Quercus petraea* in this study) which is not always feasible, while *validation using common garden data* can be used also with scarce common data networks (as is the case of *Abies alba* in this study).

### Phenotypic data

#### National Forest Inventories (NFI)

Observation data comprised ten annual campaigns of the French NFI (2005–2014; http://inventaire-forestier.ign.fr), which consists of a regular grid (1 km^2^) of temporary forest plots of 707 m^2^ each. In this study, we focused on French NFI (Appendix S1, Fig. S1.1) because inventories of neighbouring countries do not provide age data, thereby preventing the effect of age to be accounted for in models. Nevertheless, the distribution of French NFI plots has a good representativeness of the climatic range of the two species (Fig. S2.1). In each NFI plot, we selected trees for which height (in m), diameter at breast height (dbh; in cm) and age (years) data were measured. In particular, tree age was estimated from wood increment cores collected at breast height (1.30 m) for the one or two of the largest dominant trees in the plot. To account for stand density and the local abundance of neighbouring trees on tree height variation among plots, we computed for each NFI plot the sum of the basal area of neighbouring trees larger than 7.5 cm dbh (Kunstler *et al.*, 2016). We removed plots outside the natural distribution range of the species (Fig. S1.1), identified as plantation or if there was any evidence of recent (<5 years) management, for example logging. We assumed that trees in the remaining plots originated from local provenances within the same bioclimatic region. The final dataset consisted of 5376 trees from 3614 plots for *Q. petraea*, and 1304 trees from 904 plots for *A. alba*.

#### Common gardens

Common garden data were used to independently validate *in-situ* models (NFI calibration). They were established for breeding purposes during 1990–1996 for *Q. petraea* and 1967–1972 for *A. alba*, as follows: (i) seeds were collected from seed sources (hereafter provenances) throughout the natural distribution range of the species (*N* = 141 for *Q. petraea, N* = 47 for *A. alba);* (ii) the seeds were sown in a nursery; (iii) seedlings were transplanted to several sites, i.e., common gardens (N = 13 for *Q. petraea*, *N* = 6 for *A. alba*; Fig. S1.1), using a randomised block design; and (iv) measurements of tree height were made at several different years. To avoid pseudo-replication, we randomly selected a single measurement year for each tree. Neighbour basal area was assumed to be constant because plantations have a regular spacing design. Tree age at the time of height measurement was considered to be the time since sowing. A detailed description of the *Q. petraea* provenance tests is provided in a previous study (Sáenz-Romero *et al.*, 2017). A description of the *A. alba* provenances studied in common gardens is provided in Appendix S1 (Tables S1-S2).

### Climate data

To analyse phenotypic trait response to long-term climate and recent climate change, we used the yearly climate grids (1901–2014) at 30 arc sec resolution (~1 km^2^) of the EuMedClim dataset covering Europe and the Mediterranean Basin (Fréjaville & Benito Garzón, 2018). For the present study, the following bioclimatic variables were considered (Fig. S2.1): annual mean temperature, maximum temperature of the warmest month, minimum temperature of the coldest month, annual precipitation, precipitation of the wettest and the driest month, annual potential evapotranspiration, potential evapotranspiration of the warmest and the coldest month and water balance (precipitation minus potential evapotranspiration) of the wettest and the driest month. EuMedClim was computed following an anomaly approach using the fine 30’ resolution of WorldClim climate means (version 1.4, Hijmans *et al.*, 2005) to adjust the coarse spatial 0.5° resolution of yearly climate data from the Climate Research Unit (version ts3.23, Harris *et al.*, 2014). EuMedClim provides inter-annual variation of bioclimatic conditions at high spatial resolution, allowing the analysis of climate at different spatial and temporal scales instead of using climate means over a reference period (e.g. WorldClim).

To fulfil the requirements of our modelling approach based on the different climate scales at which phenotypic plasticity and local adaptation act, we split the climate data into three different sets of data: (i) long-term climate (LTC) that is the average climate value of the 1901–1960 period, and represents the climate driven local adaptation in the past for provenances (for common gardens) or for a common bioclimatic origin (for NFI, assuming that that all trees have a local origin in a given bioclimatic origin – Appendix S2); (ii) short-term climate (STC) represents the plastic response of trees to recent climate and is calculated as the local climate averaged over the 10 years preceding the measurements (NFI and common garden data); iii) recent climate change (RCC), calculated by subtracting LTC from STC to avoid collinearity problems between LTC and STC in NFI.

### Models of intra-specific trait variability

Hereafter, we refer to models calibrated using NFI data as *in-situ* models and to models calibrated using common gardens as *ex-situ* models. For a given species, the phenotypic trait *T_ijk_* (tree height) of the *i*^th^ tree individual of the *j*^th^ bioclimatic region (or provenance) in the *k*^th^ plot (or common garden) was modelled as follows:

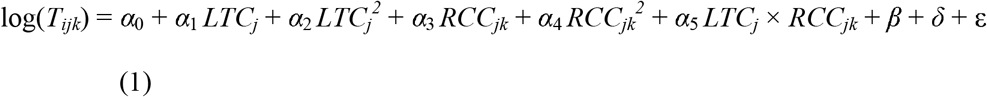

where *LTC_j_* is the long-term climate of either the *j*^th^ bioclimatic region in NFI or the *j*^th^ provenance in common gardens; *RCC_jk_* is the recent climate change defined as the difference between the STC at the *k*^th^ site (i.e., the NFI plot or the common garden) and *LTC_j_*. We included quadratic terms for both *LTC_j_* and *RCC_jk_* to consider linear and hump-shaped height responses to climate across species ranges. *β* includes ontogeny and neighbour basal area covariates and is defined as:

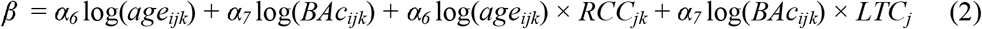

where *age* is the tree age (in years, estimated at breast height in NFI and as the time since sowing in common gardens) and *BAc* is the sum of the basal area of neighbouring trees (assumed to be constant in common gardens). *δ* gathers random effects and ε is the model error. To control for differences among sampling units in soil fertility, management (or disturbances) and environmental factors not accounted for by fixed effects, we set as random effects the plot nested within the bioclimatic region in *in-situ* models and the block nested within the site in *ex-situ* models (randomised block design). In the case of *A. alba*, the bioclimatic region random effect was not retained because it inflated p-values of LTC terms in the *in-situ* model (high redundancy). One main difference between NFI and common garden data is the age of trees (Fig. S4.1). We reduced this difference by excluding old trees (> 200 years) in NFI and saplings (< 10 years) in common gardens, and we added *age* as a covariate in the models to control for ontogeny. To control for potential differences in growth response to climate change among ontogenic stages, we added the interaction term *age_ijk_* × *RCC_jk_* in both models. We also introduced *BAc* as covariate to control for neighbour basal area (Fig. S4.1) and the interaction term *BAc_ijk_* × *LTC_j_* to control for potential differences in neighbour basal area effects among bioclimatic regions in *in-situ* models. A saturated model form including *BAc_ijk_* × *RCC_jk_* and *age_ijk_* × *LTC_j_* interaction terms was not retained as it decreased model parsimony and the significance of parameters of interest. Models were fitted using the R package nlme (Pinheiro *et al.*, 2015). Coefficients of determination were used to compute the percentage of explained variance by fixed effects alone (R^2^_marginai_) and both fixed and random effects (R^2^_conditional_) (Nakagawa & Schielzeth, 2013).

For each species, we selected one single explanatory bioclimatic variable to represent LTC and RCC, the same between the two datasets (to enable comparison). The variable selection process was as follows. First, we fitted one model per dataset for each bioclimatic variable (Fig. S2.1) using eqns (1-2). Second, we removed models when parameter estimates for LTC and RCC were not significant at *P* = 0.1 or when positive quadratic relationships were fit (*a_2_* > 0 or *α_4_* > 0) to keep models with decreasing tree height towards one or both ends of the climatic gradient. Third, competitive models were compared using the Akaike information criterion (AIC), and the final model selection was based on the lowest AIC values for both *in-situ* and *ex-situ* models (to enable comparison).

### Model predictions

#### Separating local adaptation, plasticity and their interaction

Model coefficients were used to separate components of phenotypic variation by substituting RCC to its climatic components (*RCC_jk_* = *STC_k_* - *LTC_j_*) in eqn. (1):

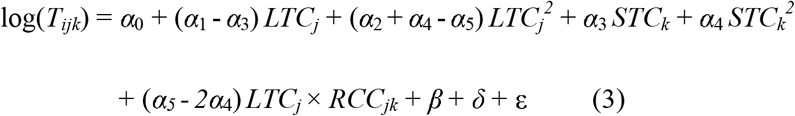

This analytical decomposition enables to estimate the relative effects of the long-term and short-term climate in the field, using RCC and LTC from eqn. (1). From eqn. (3), coefficients associated to linear (*α*_1_ − *α*_3_) and quadratic ((*α*_2_ + *α*_4_ − *α*_5_) variation of LTC are used to predict the effect of the provenance (local adaptation), coefficients associated to linear (*α*_3_) and quadratic (*α*_4_) variation of STC are used to predict phenotypic plasticity (reaction norms) and those associated to *LTC_j_* × *STC_k_*(*α*_5_ − *2α*_4_) are used to predict their interaction.

#### Spatial predictions

The *in-situ* and *ex-situ* models were used to make spatial predictions of provenance and plasticity effects, their interaction and the total component of tree height response to climate across Europe. We computed *LTC_j_* and *STC_k_* in each grid cell (30 arc sec resolution) respectively using the long-term (1901–1960) and the recent short-term (2001–2014) averaged values of the corresponding climatic variables to compute maps according to fitted parameters in eqn. (3). Effects of covariates were fixed for *age* (12-year-old trees), *BAc* (30 m^2^ ha^−1^) and for their interaction with *RCC_jk_* and *LTC_j_* (both averaged across the species natural range), respectively, according to eqn. (2), and these constants (including the intercept *α*_0_) were added to the total variation component.

### Model validation

To validate our approach, we used two alternative methods (Fig. 1) to test the ability of *in-situ* models to predict independent tree height observations in common gardens where the effects of the provenance, plasticity and their interaction can be separated.

#### Validation using common garden data

To compare the predictions of *in-situ* models with raw common garden data, we first predicted the mean height of each provenance in each site (provenance-by-site mean) from eqn. (3), as a function of the LTC of the provenance and the STC of the site for a given age and neighbour basal area. Then, we compared these predictions to observed values in common gardens. Both predicted and observed provenance-by-site means were standardized across sites and provenances to estimate provenance and plasticity effects, respectively (see Appendix S3). Correlations between predicted and observed values were tested using Pearson correlation coefficients.

#### Validation using ex-situ model predictions

To compare the predictions of *in-situ* and *ex-situ* models, we predicted provenance and plasticity effects, and their interaction, as a function of LTC and STC conditions in common gardens using both *in-situ* and *ex-situ* models (see Appendix S3). From eqn. (3), the mean tree height of a provenance planted in several common gardens was predicted for a given age and neighbour basal area as a function of the LTC of the provenance (genetic effect) and the mean tree height of each provenance was predicted as a function of the STC of the site (plasticity of the provenance). Correlations between paired predictions from *in-situ* and *ex-situ* models of provenance and plasticity effects, their interaction and the sum of all three components of tree height (total variation) were tested using Pearson correlation coefficients. For the interaction component, correlation coefficients were computed separately for the mean plastic responses (reaction norms) of cold, core and warm provenances. We classified provenances among cold, core and warm parts of the range using the 1-33^th^, 34-66^th^ and 67-100^th^ percentiles of LTC, respectively, computed across the natural distribution range of the species.

## RESULTS

We found both *in-situ* and *ex-situ* models with significant terms for LTC, RCC and their interaction on individual tree height in *Q. petraea* whereas only the *in-situ* model was found significant for *A. alba*. In *Q. petraea*, selected *in-situ* and *ex-situ* models were based on the maximum temperature of the warmest month (Tmax). In *A. alba*, the selected *in-situ* model was based on the potential evapotranspiration of the warmest month (PETmax). Both *in-situ* and *ex-situ* models indicated significant negative interaction between LTC and RCC (Table 1) and positive interaction between LTC and STC (Table 2) in both species. Hence, average plasticity differed among regions and provenances for both species.

**Table 1.** Estimates of *in-situ* and *ex-situ* linear mixed-effect models of individual tree height data (log-transformed). *In-situ* models were fit on 5376 trees (height measurements) in 3617 plots nested in **22** bioclimatic regions (random groups) for *Q. petraea* and on 1304 trees in 904 plots nested in 15 bioclimatic regions for *A. alba*. Bioclimatic regions were added as random effects in *in-situ* model for *Q. petraea* but not for *A. alba*. *Ex-situ* models were fit on 130241 trees (height measurements) from 141 provenances that were planted in 591 blocs nested in 13 sites (random groups) for *Q. petraea* and on 13314 trees from 47 provenances planted in 166 blocs nested in **6** sites for *A. alba*. No significant *ex-situ* models were found in *A. alba*. The climatic variable used to compute LTC and RCC is the maximal temperature of the warmest month in *Quercus petraea* and the potential evapotranspiration of the warmest month in *Abies alba*. The percentage of the variance explained by the models is measured by the marginal (fixed effects, m) and conditional (both fixed and random effects, c) adjusted R^2^. ‘Df’ degree of freedom, ‘BAc’ sum of basal area of neighbouring competitor trees, ‘LTC’ long-term climate, ‘RCC’ recent climate change.

| *Quercus*<br>*petraea* | *in-situ* |  |  |  | *ex-situ* |  |  |  |
| --- | --- | --- | --- | --- | --- | --- | --- | --- |
|  | Estimate (±SE) | Df | t-value | *P* | Estimate (±SE) | Df | t-value | *P* |
| Intercept | -13.40 (4.07) | 3589 | -3.3 | 0.001 | -12.38 (2.23) | 129643 | -5.54 | <0.001 |
| log(age) | 0.311 (0.009) | 1758 | 34.53 | <0.001 | 1.322 (0.008) | 129643 | 167.71 | <0.001 |
| log(BAc) | -0.171 (0.133) | 1758 | -1.29 | 0.197 |  |  |  |  |
| LTC | 1.269 (0.354) | 19 | 3.59 | 0.002 | 0.903 (0.181) | 129643 | 5 | <0.001 |
| LTC<sup>2</sup> | -0.028 (0.008) | 19 | -3.58 | 0.002 | -0.020 (0.004) | 129643 | -5.55 | <0.001 |
| RCC | 0.440 (0.118) | 3589 | 3.72 | <0.001 | 0.730 (0.182) | 129643 | 4.01 | <0.001 |
| RCC<sup>2</sup> | -0.017 (0.003) | 3589 | -5.22 | <0.001 | -0.018 (0.004) | 129643 | -4.95 | <0.001 |
| LTC:RCC | -0.018 (0.005) | 3589 | -4.02 | <0.001 | -0.031 (0.007) | 129643 | -4.26 | <0.001 |
| log(BAc):LTC | 0.013 (0.006) | 1758 | 2.38 | 0.017 |  |  |  |  |
| log(age):RCC | 0.009 (0.007) | 1758 | 1.3 | 0.193 | -0.012 (0.003) | 129643 | -3.63 | <0.001 |
| R2 (m/c) (%) | 41/91 |  |  |  | 26/65 |  |  |  |
| *Abies*<br>*alba* | Estimate (±SE) | Df | t-value | *P* |  |  |  |  |
| Intercept | 0.732 (0.649) | 898 | -1.13 | 0.259 |  |  |  |  |
| log(age) | 0.319 (0.014) | 396 | 23.57 | <0.001 |  |  |  |  |
| log(BAc) | 0.224 (0.123) | 396 | 1.82 | 0.069 |  |  |  |  |
| LTC | 0.024 (0.008) | 898 | 2.95 | 0.003 |  |  |  |  |
| LTC<sup>2</sup> | -7.3 10<sup>-5</sup> (2.8 10<sup>-5</sup>) | 898 | -2.63 | 0.009 |  |  |  |  |
| RCC | 0.009 (0.006) | 898 | 1.46 | 0.146 |  |  |  |  |
| RCC<sup>2</sup> | -9.9 10<sup>-5</sup> (2.3 10<sup>-5</sup>) | 898 | -4.36 | <0.001 |  |  |  |  |
| LTC:RCC | -8.3 10<sup>-5</sup> (3.8 10<sup>-5</sup>) | 898 | -2.17 | 0.030 |  |  |  |  |
| log(BAc):LTC | -2.4 10<sup>-4</sup> (9.5 10<sup>-4</sup>) | 396 | -0.25 | 0.803 |  |  |  |  |
| log(age):RCC | 0.002 (8.4 10<sup>-4</sup>) | 396 | 1.92 | 0.056 |  |  |  |  |
| R2 (m/c) (%) | 49/86 |  |  |  |  |  |  |  |

**Table 2.** Mean bootstrap estimates (± SD) of tree height variation due to provenance and plasticity effects, and their interaction, computed from *in-situ* (NFI) model and *ex-situ*(common garden) model. ‘x’ and ‘x^2^’ indicate linear and quadratic terms respectively. Significant differences of bootstrapped values (n = 200) to null values were tested using t-tests; all are significant at *P* < 0.001.

|  | Provenance effect |  | Phenotypic plasticity |  | interaction |
| --- | --- | --- | --- | --- | --- |
|  | x | x<sup>2</sup> | x | x<sup>2</sup> |  |
| *Quercus petraea*<br>*ex-situ* model | 0.191<br>(0.026) | -7.3 10<sup>-3</sup><br>(0.35 10<sup>-3</sup>) | 0.712<br>(0.169) | -0.017<br>(0.003) | 4.6 10<sup>-3</sup><br>(0.68 10<sup>-3</sup>) |
| *Quercus petraea*<br>*in-situ* model | 0.548<br>(0.365) | -0.022<br>(0.009) | 0.465<br>(0.122) | -0.018<br>(0.003) | 0.018<br>(0.007) |
| *Abies alba*<br>*in-situ* model | 0.014<br>(0.007) | -8.6 10<sup>-5</sup><br>(2.1 10<sup>-5</sup>) | 0.010<br>(0.007) | -9.2 10<sup>-5</sup><br>(2.1 10<sup>-5</sup>) | 1.0 10<sup>-4</sup><br>(0.4 10<sup>-4</sup>) |

*In-situ* selected models indicated significant positive effects of tree age and neighbour basal area on tree height (Table 1, Fig. S4.1). Tree height increased with increasing neighbour basal area in *A. alba* (*P* = 0.07) and with neighbour basal area towards warmer regions in *Q. petraea* (positive interaction between BAc and LTC, *P* = 0.02; Table 1).

### Statistical approximation of provenance and plasticity effects

#### Sessile oak (Quercus petraea)

Validation using common garden data indicated that predictions of tree height variation among provenances and among sites using the *in-situ* model were significantly correlated with observations in common gardens in *Q. petraea* (*P* < 0.001 for estimates of provenance and plasticity effects, and the total component of tree height, Fig. S6.3). Validation using *ex-situ* model predictions showed high correlations between *in-situ* and *ex-situ* model predictions of the provenance and plasticity effects, and their interaction (Table 2, Fig. 2). We found significant quadratic responses of height to Tmax for the provenance effect (Fig. 2a), plasticity (Fig. 2b) and the total (Fig. 2c) variation, that were similar between *in-situ* and *ex-situ* models with high correlations between paired predictions (*r* > 0.80, *P* < 0.001). We found high correlations between *ex-situ* and *in-situ* model paired predictions of cold (*r* = 0.65, *P* < 0.001), core (*r* = 0.99, *P* < 0.001) and warm provenance mean reaction norms (*r* = 0.97, *P* < 0.001). Both models showed similar patterns in Tmax optimums (i.e. Tmax values corresponding to maximum predicted heights) among cold, core and warm provenances that were respectively warmer and colder for warm and cold provenances (Kruskal-Wallis tests, *P* < 0.001, Fig. 2d-e). The *in-situ* model, however, showed higher differences in optimums among provenances (Fig. 2e).

**Figure 2.**
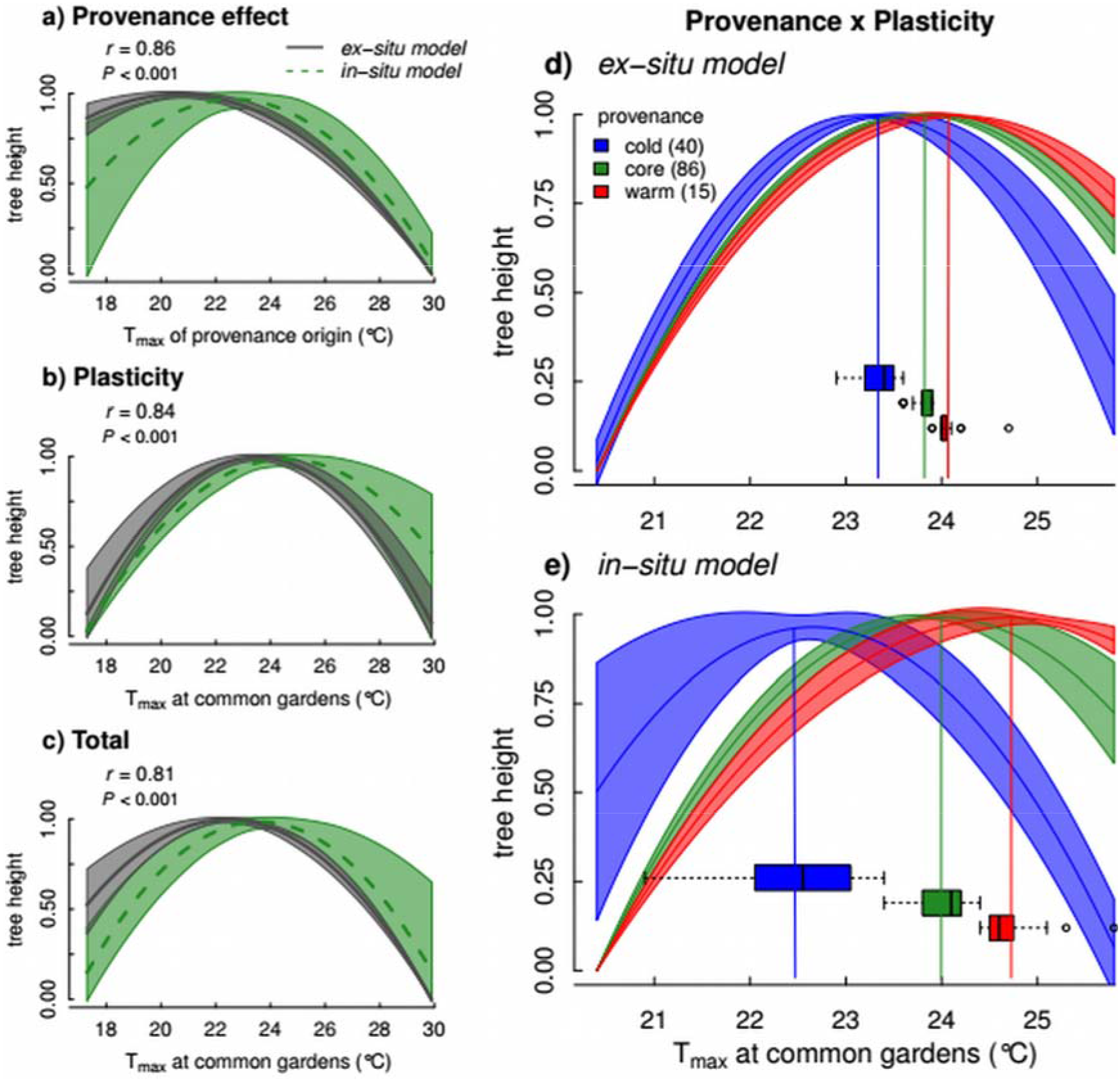
Comparison of *in-situ* model (NFI) and *ex-situ* model (common gardens) predictions of provenance (a) and plasticity effects (b), and the total component (c) of tree height variation, recorded in common gardens in *Quercus petraea*. Pearson correlation coefficients between *ex-situ* and *in-situ* model predictions are reported. (d-e) Model predictions of plastic responses among provenances (provenance × plasticity interaction). Temperature optima for cold, core and warm provenances are indicated by horizontal boxplots; vertical coloured lines indicate mean optimum values. Significant differences in temperature optimum were tested using Kruskal-Wallis tests: χ^2^= 105.2 in d) and χ^2^ = 104.2 in e), *P* < 0.001 for both. Shaded areas and lines represent the standard deviation around average model predictions (computed by bootstrapping in a-c). Predictions were scaled between 0–1 independently for *in-situ* and *ex-situ* models. Model parameters (coefficients and significance) are presented in Table 1. Validation analyses using common garden data are presented in Appendix S3.

#### Silver fir (Abies alba)

In *A. alba*, the *in-situ* model showed significant quadratic responses of tree height to PETmax for the provenance and plasticity effects (Table 2). Validation using common garden data indicated that the *in-situ* model significantly predicted tree height variation among provenances in common gardens (provenance effect: *r* = 0.44, *P* < 0.01, Fig. 3a) and among sites (plasticity effect: *r* = 0.72, *P* < 0.001, Fig. 3b). The correlation was weak for the total variation (*r* = 0.20, *P* = 0.06, Fig. 3c, Fig. S7.3). The *in-situ* model predicted warmer optimums for warm provenances and colder optimums for cold provenances (Kruskal-Wallis tests, *P* < 0.001, Fig. 3d).

**Figure 3.**
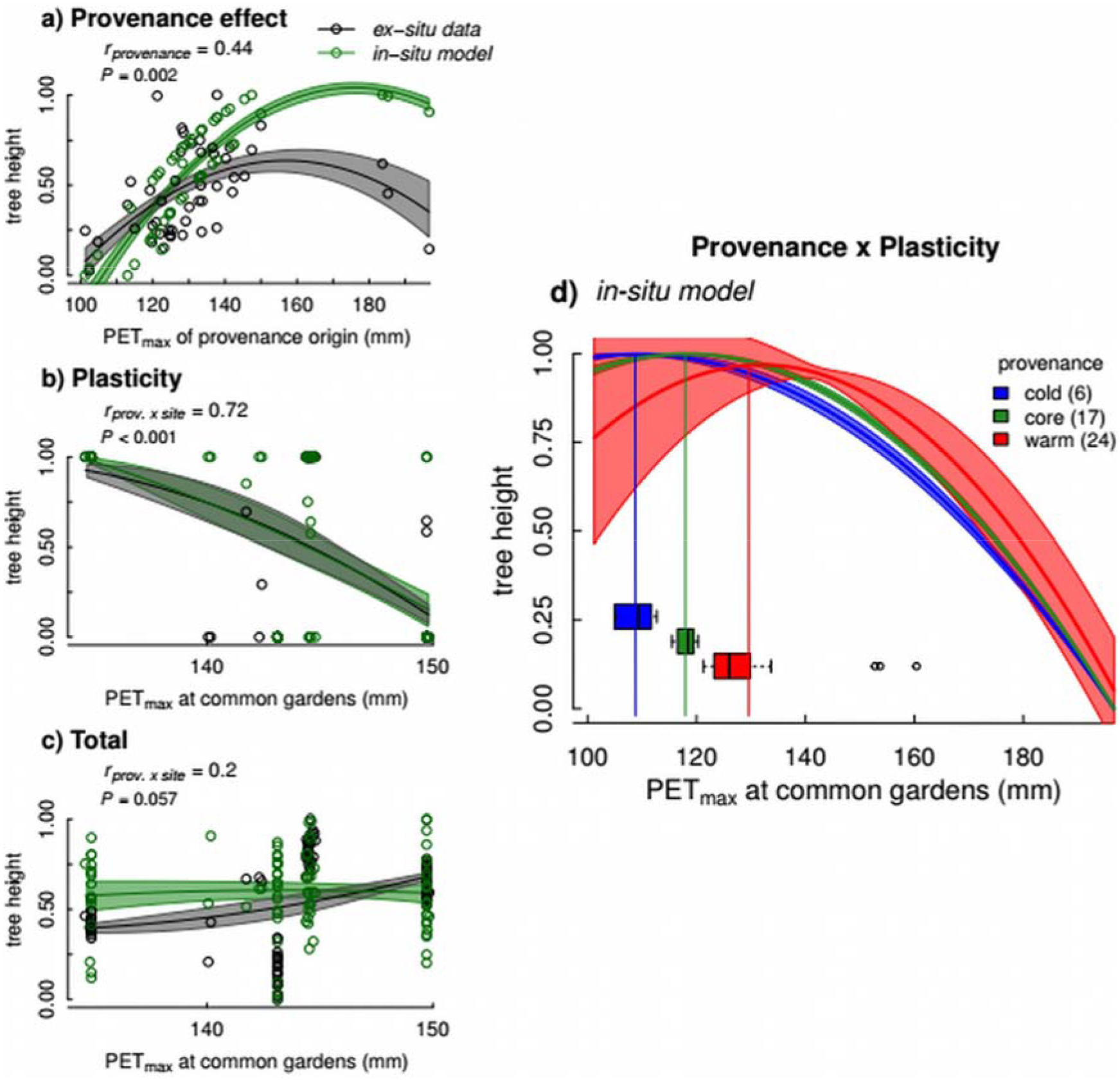
Comparison of *in-situ* (NFI) model predictions and common garden data (*ex-situ*) estimates of provenance (a) and plasticity effects (b), and the total component (c) of tree height variation, recorded in common gardens in *Abies alba*. Points represent provenance means in a) and provenance-by-site means in b-c that were computed according to the validation method using common garden data (see Fig. 1). Computation of provenance and plasticity effects, and the total variation is described in Appendix S3. Pearson correlation coefficients between *ex-situ* data and *in-situ* model predictions are reported. (d) *In-situ* model predictions of plastic responses among provenances (provenance × plasticity interaction). Temperature optima for cold, core and warm provenances are indicated by horizontal boxplots; vertical coloured lines indicate mean optimum values. Significant differences in temperature optimum were tested using Kruskal-Wallis tests: χ^2^ = 105.2, *P* < 0.001. Shaded areas and lines represent the standard deviation around average model predictions (computed by bootstrapping in a-c). Height values were scaled between 0–1 independently for *in-situ* predictions and common garden data. *In-situ* model parameters (coefficients and significance) are presented in Table 1.

### Range-wide spatial predictions of tree height

In *Q. petraea*, both the *ex-situ* and *in-situ* models predicted very similar spatial patterns (Fig. 4) in the relative variation of tree height for provenance and plasticity effects, and their interaction, despite differences in absolute values for total variation for a given age and neighbour basal area (Fig. 4g-h). These differences are explained by the differences in tree age estimation between the two datasets (i.e. age is measured at breast height in NFI and as the time since sowing in common gardens). The quadratic response of tree height to LTC (Fig. 2a) predicted that trees living at the warm limit of the species range were the shortest, which is illustrated by the short heights predicted over southern Europe from the provenance effect (transparent colours, Fig. 4a-b). Similarly, trees inhabiting the warmest conditions in the southernmost part of the species distribution range were also predicted to be shorter from the phenotypic plasticity effect (Fig. 4c-d). Spatial predictions of interaction effects showed an opposite pattern with increasing height towards warmer climates (Fig. 4e-f). Total variation followed spatial patterns of the provenance and plasticity effects (Fig. 4g-h). In *A. alba*, trees inhabiting cold climates (e.g. high elevation areas in the Alps) were predicted to be shorter according to the *in-situ* model (Fig. 5). In warm climates, the provenance effect (Fig. 4a) and provenance × plasticity interaction (Fig. 5c) predicted taller trees while plasticity predicted smaller trees over southern Europe (Fig. 5b).

**Figure 4.**
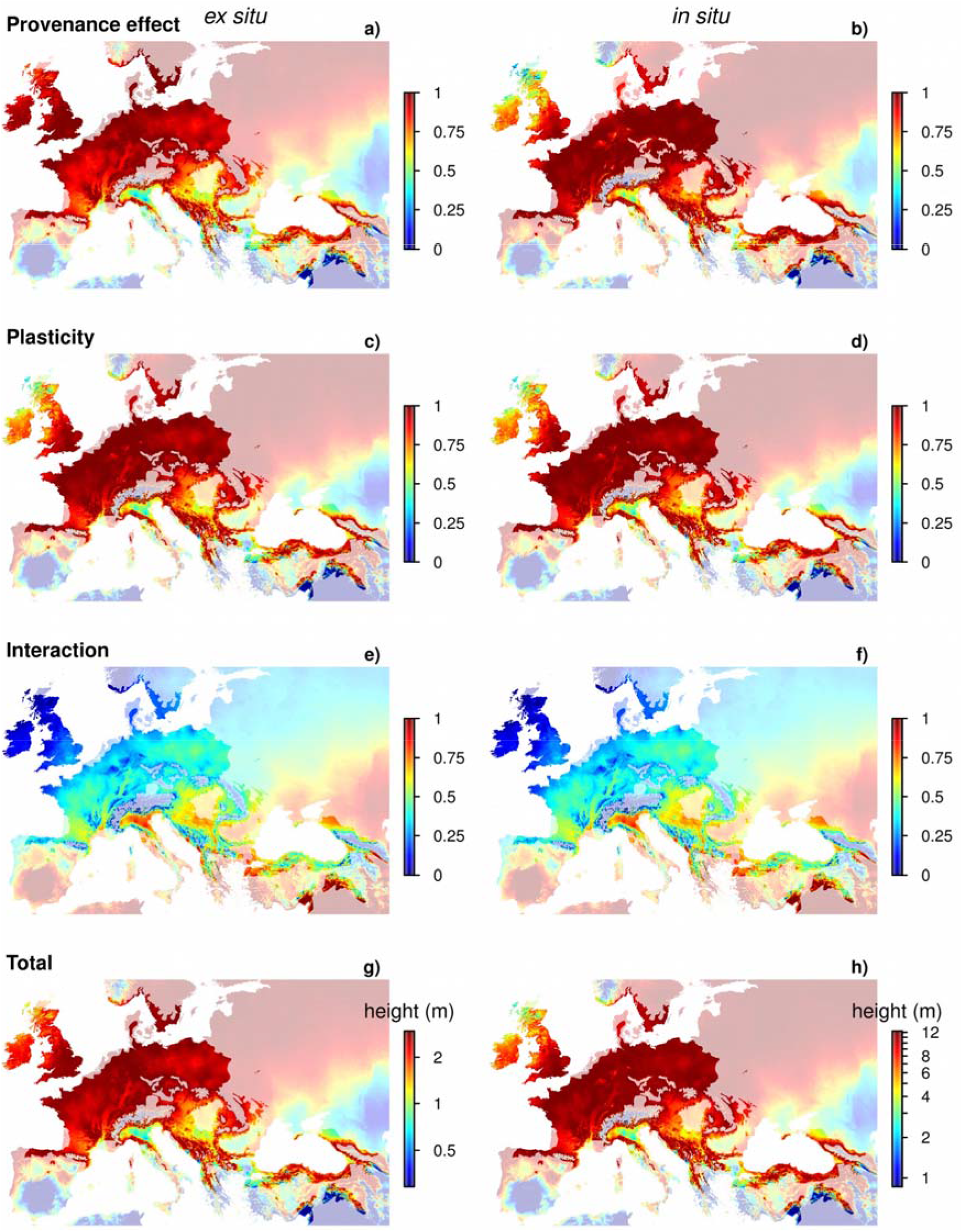
Spatial predictions of *Quercus petraea* range-wide variation in tree height using *ex-situ* (a, c, e, g) and *in-situ* models (b, d, f, h). Maps indicate the provenance effect (a-b), plasticity (c-d), their interaction (e-f) and the total variation of tree height (g-h). The shaded area represents model predictions outside the natural distribution range of the species. Predictions are for 12-years-old trees, with neighbour basal area set to average conditions (30 m^2^ ha^−1^) in the *in-situ* model.

**Figure 5.**
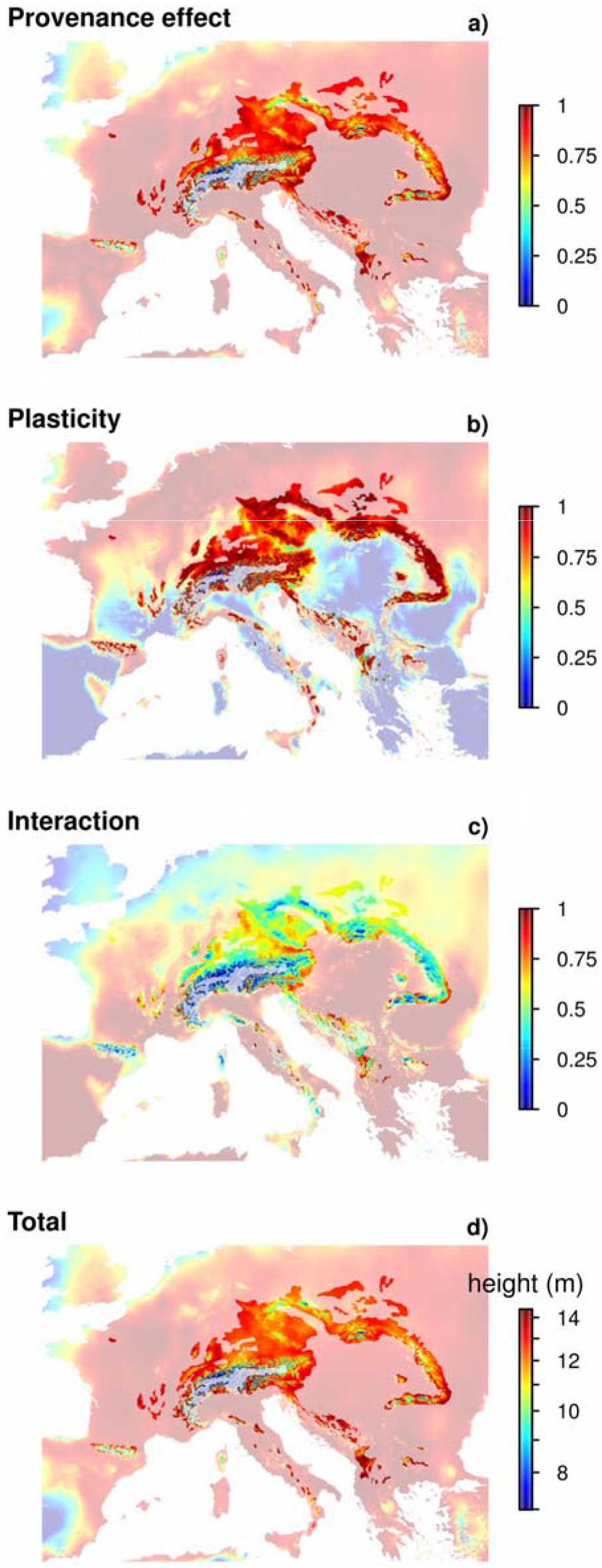
Spatial predictions of *Abies alba* range-wide variation in tree height using the *in-situ* model. Maps indicate the provenance effect (a), plasticity (b), their interaction (c) and the total variation of tree height (d). The shaded area represents model predictions outside the natural distribution range of the species. Predictions are for 12-years-old trees, with neighbour basal area set to average conditions (30 m^2^ ha^−1^).

## DISCUSSION

### Can local adaptation and phenotypic plasticity in tree height be inferred from *in-situ*observations?

To date, disentangling the sources of phenotypic variation has only been addressed by analysing common gardens or reciprocal transplant experiments (e.g. Kawecki & Ebert, 2004; Hoffmann & Sgrò, 2011; Blanquart *et al.*, 2013; Latreille & Pichot, 2017), using similar approaches as those that we named here *ex-situ* models. However, as common garden data are often scarce, we propose an alternative method for understanding the causes of phenotypic variation: using increasingly abundant data from field observations, such as NFI. Using the rationale of *ex-situ* models based on common garden data, we defined *in-situ* models based on NFI data and thoroughly validated *in-situ* model predictions with raw data coming from common gardens and with predictions from *ex-situ* models.

Overall, our results show that *in-situ* models correctly predicted phenotypic patterns observed in common gardens (Table 2, Figs 2-5), suggesting that field observations (NFI) can be used to statistically approximate the range-wide intraspecific variation in tree height that is attributable to local adaptation (provenance effects), plasticity and their interaction components. In particular, our results suggest that differences among provenances that are related to their climate of origin can be statistically approximated using field measurements by modelling trait variation as a function of the long-term regional climate (LTC), while the recent climate change (RCC) and its components (STC - LTC) can be used to estimate the plastic response of the trait. However, we must keep in mind that other models can achieve similar predictions. Hence further studies are needed to decipher whether our approach can be used to estimate local adaptation and phenotypic plasticity components of intraspecific variation in the field for other traits and taxa.

In both species, the most parsimonious models (both *in-situ* and *ex-situ)* were based on climatic variables related to summer temperature: PETmax in *A. alba* and Tmax in *Q. petraea*. This underlies the high sensitivity of *Abies alba* to the evaporative demand in summer (Lebourgeois *et al.*, 2013) and that temperature chiefly drove differences in tree height among *Q. petraea* provenances while drought mostly drove plastic responses for tree height and survival in this species (Sáenz-Romero *et al.*, 2017).

Although validation analyses showed good agreement between *in-situ* model predictions and common garden data for both species, we found some discrepancies. In *A. alba*, we did not find any significant *ex-situ* model. This may be explained by the fact that less common garden data are available for this species compared to *Q. petraea*. In addition, *A. alba* is a generalist species with limited local adaptation to temperature and water availability (Frank *et al.*, 2017; Latreille & Pichot, 2017). Otherwise, predictions based on NFI data in *Q. petraea* (Fig. 2d) showed higher differences of climatic optima among provenances than those based on common garden data (Fig. 2e). These differences might be induced by intrinsic differences in the nature of data. The large difference in the age of trees (older in NFI data) and the fact that NFI data rely on dominant trees might partly explained these differences and the fact that a stronger signal of local adaptation (i.e. higher differences in climatic optima) was found using NFI data.

### Common patterns of local adaptation and plasticity in tree height: implications for species distribution ranges under climate change

In both species, we found hump-shaped relationships between height and climate for the provenance (long-term climate) and plasticity effects (short-term climate) and a positive interaction effect between them (Table 2, Figs 2-3). The latter indicated that climatic optima of provenances co-vary positively with their climate of origin: warmer provenances grow taller in warmer climates and colder provenances grow taller in colder climates (Figs 2d-e and 3d). In addition, this indicated that plasticity differs significantly among populations, as for other species (Wang *et al.*, 2006; Leites *et al.*, 2012; Münzbergová *et al.*, 2017; Sáenz-Romero *et al.*, 2017). This consistency between *A. alba* (a mountain evergreen conifer tree) and *Q. petraea* (a temperate deciduous broadleaved tree) points to potential common patterns in local adaptation and plasticity among tree species, as recently indicated in boreal conifer trees (Pedlar & McKenney, 2017).

Our spatial predictions of phenotypic plasticity suggest that tree height of warm provenances has decreased in response to recent climate warming, mostly in southern Europe (Figs 4-5). These results suggest that recent warming may have pushed species at the warmest boundary of the distribution range beyond their tolerance limits, that is corroborated by a higher mortality in warmest/driest range margins for *A. alba, Q. petraea* and other European tree species (Cailleret *et al.*, 2014; Benito Garzón *et al.*, 2018). Furthermore, we found that the effect of neighbour basal area on tree height was dependent on the climate of the bioclimatic region in *Q. petraea*, emphasising that tree sensitivity to biotic interaction (e.g. competition) may change along climatic gradients (Gomez-Aparicio *et al.*, 2011; Kunstler *et al.*, 2016).

## CONCLUSION

We show that local adaptation and plasticity components of phenotypic variation, and their interaction, can be statistically approximated using field observations of wild tree populations subject to recent climate warming. However, further studies are needed to determine whether the ability of *in-situ* models to predict trends in common garden experiments represent a shared underlying cause that can be generalized to other situations, i.e. whether climate variations at different scales can be used to separate local adaptation and plastic responses to climate in field conditions. The modelling framework used in our study could be applied to many species and traits, offering a promising avenue to enhance our understanding of local adaptation and plasticity patterns across large geographical gradients.

## DATA ACCESSIBILITY

Raw data can be freely accessed online for French National Forest Inventories (http://inventaire-forestier.ign.fr), *Quercus petraea* (https://arachne.pierroton.inra.fr/QuercusPortal/) and *Abies alba* common gardens (online repository, under process). Climate data used for this study are available online (http://gentree.data.inra.fr/climate/). R codes used for data analyses can be obtained from the correspondence author upon request.

## BIOSKETCH

The authors’ research aims to understand how ecological and evolutionary processes drive the effects of global changes on forests, by merging expertise in ecological modelling, genetics and conservation.

T.F. and M.B.G. conceived and designed the study; T.F. conducted the data analyses; T.F. and M.B.G. wrote the manuscript that was commented and improved by B.F., A.K. and A.D.

## Acknowledgements

This study was funded by the “Investments for the Future” program IdEx Bordeaux (ANR-10-IDEX-03-02) and the European Union’s Horizon 2020 research and innovation programme project GenTree (grant agreement No 676876). We are very grateful to Juan Fernandez Manjarrés (CNRS-Université Paris-Sud) with whom the discussion about how to use NFI for splitting genetic and plastic effects started many years ago. We are indebted to Denis Vauthier and Franck Rei (INRA UEFM, Avignon), Fabrice Bonne, Thierry Paul and Vincent Rousselet (INRA UEFL, Nancy) and Jean Gauvin (INRA UGBFOR, Orléans) for data collection in the *Abies alba* provenance tests. We acknowledge colleagues from different European institutions for providing access and use of provenance data in *Quercus petraea*: Jon Kehlet Hansen, University of Copenhagen, for Denmark; Brigitte Musch and the staff of the French Office National des Forêts for France; Hans-Martin Rau, Jochen Kleinschmit, Josef Svolba of Niedersächsische Forstliche Versuchsanstalt, Abteilung Forstpflanzenzüchtung, Staufenberg, Escherode and Wilfried Steiner, Alwin Janssen, Nordwestdeutsche Forstliche Versuchsanstalt, Abteilung Waldgenressourcen, Hann-Münden, for Germany; Wladyslaw Chalupka, Henryk Fober of the Instytut Dendrologii PAN, Kórnik for Poland; Steve J Lee, Alan M. Fletcher and Edward P. Cundall, Northern Research Station, for the United Kingdom.

