## Supplementary Information for "Inferring phenotypic plasticity and local adaptation to climate across tree species ranges using forest inventory data"

**Appendix S1 Phenotypic and climate data**

**Fig. S1.1** Maps of *NFI* and *common garden* tree height data for *Quercus petraea* and *Abies alba*.

**Fig. S2.1** Principal component Analysis of climatic conditions across *NFI* and *common garden* data for *Quercus petraea* and *Abies alba*.

**Fig. S3.1** Temporal trends (1901–2014) in annual mean temperature across species ranges*.*

**Fig. S4.1** Tree height variation as a function of tree age and neighbour basal area.

**Table S1** Description of the 6 common gardens used for measuring tree height on *Abies alba* provenances.

**Table S2** Description of *Abies alba* provenances planted in the 6 common gardens.

**Appendix S2 Bioclimatic regionalisation of species’ natural distribution ranges**

**Fig. S5.2** Maps of bioclimatic regions within the natural distribution range of *Quercus petraea* and *Abies alba.*

**Appendix S3 Model Validation**

*Validation using common garden data*

**Fig. S6.3** Correlation between *in-situ* model predictions and common garden data estimates of provenance and plasticity effects and the total component of variation in tree height in *Quercus petraea*.

**Fig. S7.3** Correlation between *in-situ* model predictions and common garden data estimates of provenances and plasticity effects and the total component of variation in tree height in *Abies alba*.

*Validation using ex-situ model predictions*

**Appendix S1 Phenotypic and climate data**

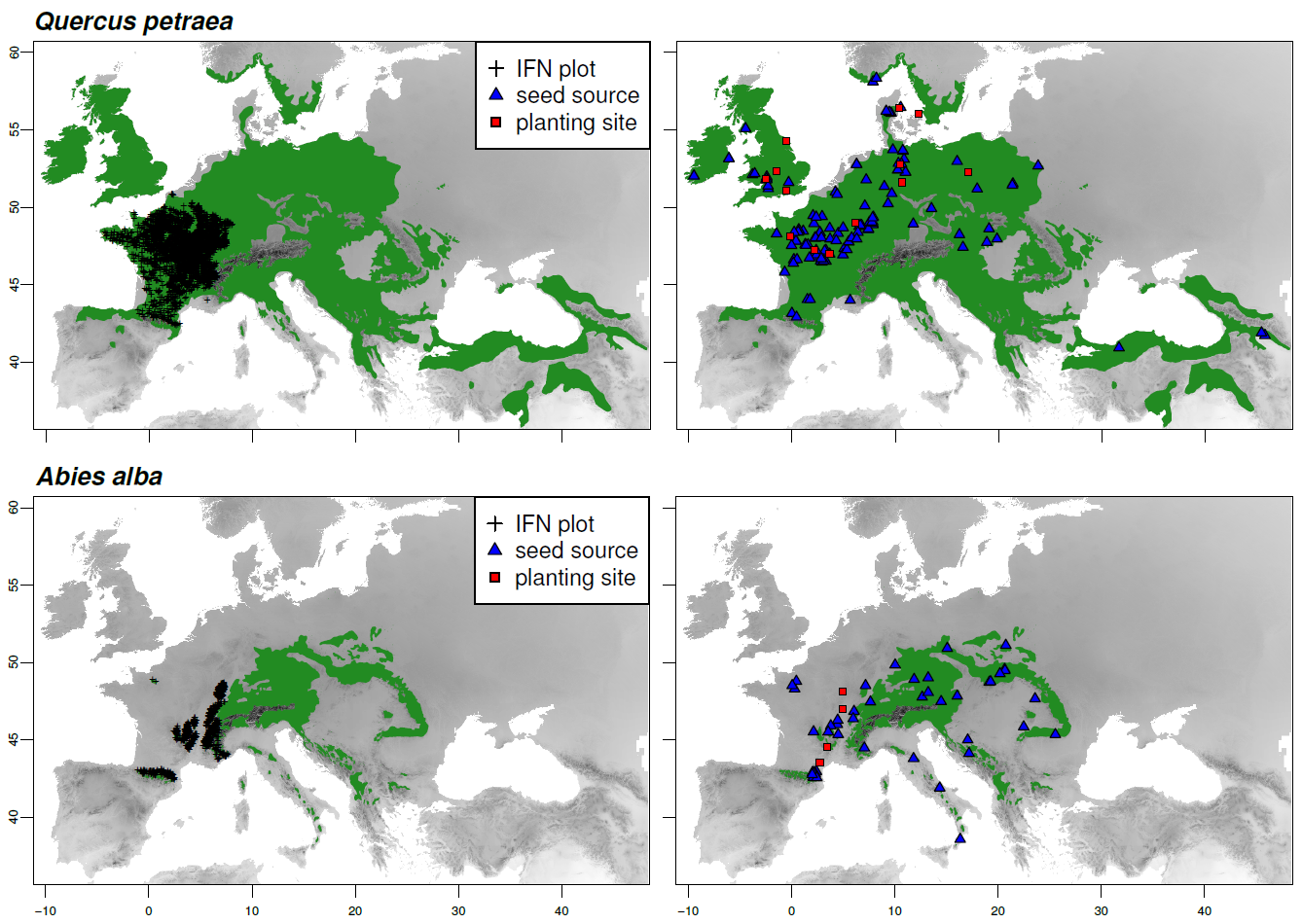

**Figure S1.1** Geographic distribution of French National Forest Inventory data (NFI) and common garden experiments within the natural distribution range of *Quercus petraea* and *Abies alba*. Black crosses represent NFI temporary forest plots (left panels) in which height, age and diameter at breast height were measured on dominant trees, within the natural distribution range of the species (green area; [http://www.euforgen.org](http://www.euforgen.org/)). Common garden experiments (right panels) consist of planted trees from provenances (seed sources, blue triangles) covering the species distribution range in common gardens (genetic trials, red squares).

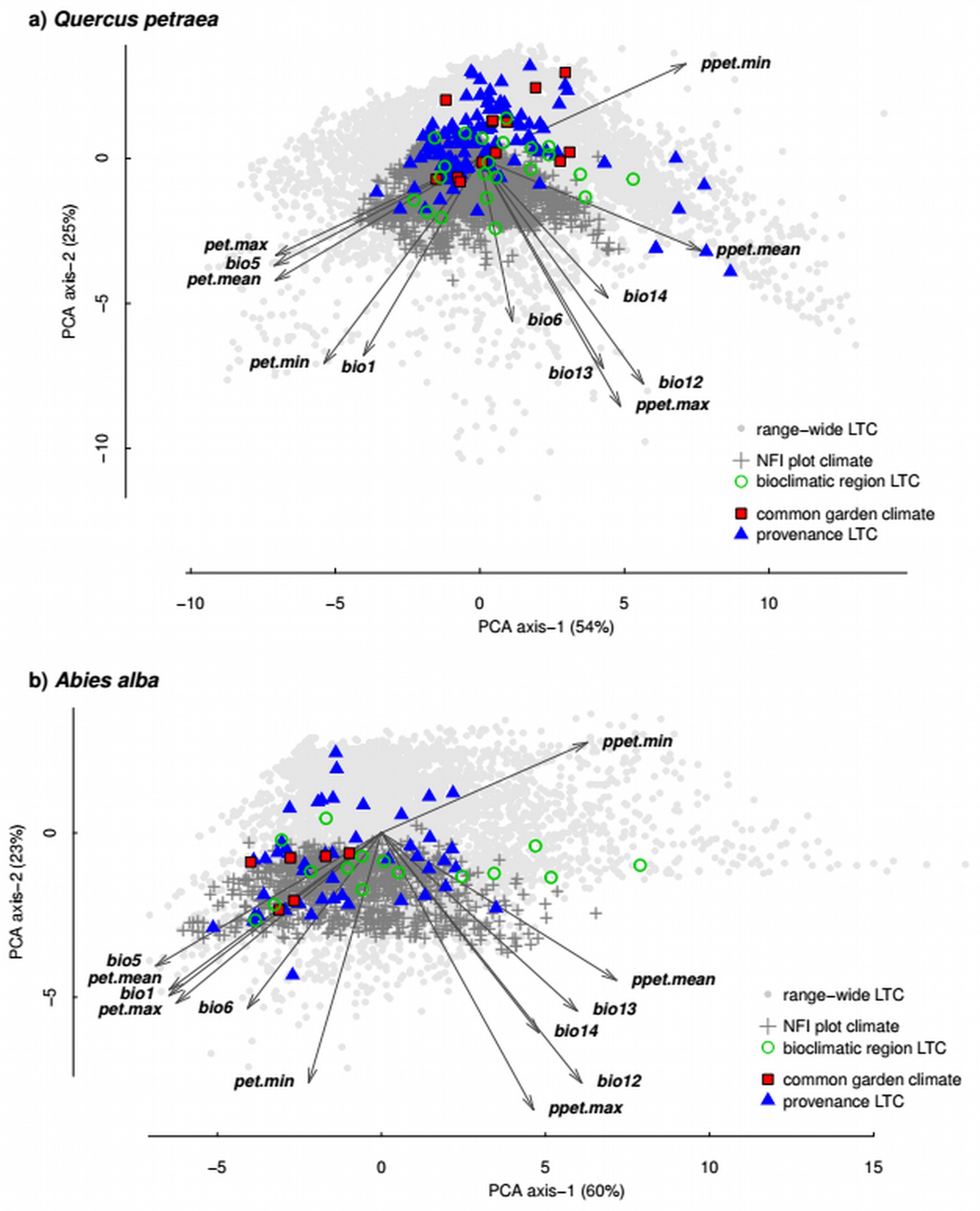

**Figure S2.1** Principal component analysis (PCA) of short-term climate at *in-situ* National Forest Inventory (NFI) plots (grey crosses) and *ex-situ* common gardens (red squares), and of long-term climate (LTC) of origin of populations planted in common gardens (‘provenance’, blue triangles) and of bioclimatic regions covered by NFI plots (black circles, see Appendix S2) for *Quercus petraea* (a) and *Abies alba* (b). The LTC distribution across the species natural distribution range is indicated by grey dots. Short-term climate represents the 10-yr means before tree measurements and LTC the 1901–1960 means. The climatic variance explained by the first two PCA axes is indicated in brackets. The PCA was computed using the following climatic variables downscaled from EuMedClim: ‘bio1’ annual mean temperature, ‘bio5’ maximum temperature of the warmest month, ‘bio6’ minimum temperature of the coldest month, ‘bio12’ annual precipitation, ‘bio13’ maximum precipitation of the wettest month, ‘bio14’ minimum precipitation of the driest month, ‘pet.mean’ annual potential evapotranspiration, ‘pet.max’ potential evapotranspiration of the warmest month, ‘pet.min’ potential evapotranspiration of the coldest month, ‘ppet.mean’ annual water balance (precipitation minus evapotranspiration), ‘ppet.max’ water balance of the wettest month, ‘ppet.min’ water balance of the driest month.

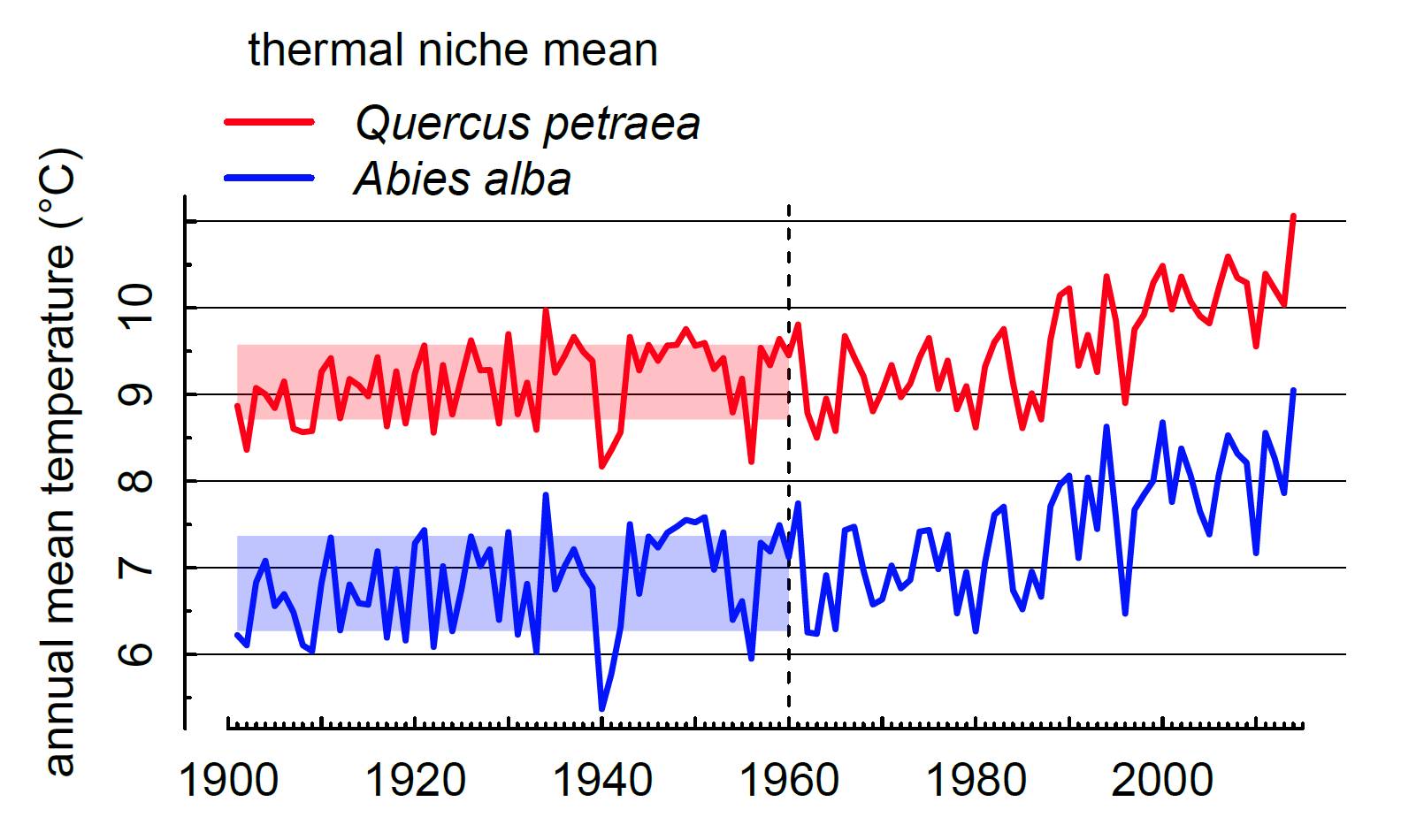

**Figure S3.1** Temporal variation (1901–2014) in annual mean temperature (from http://gentree.data.inra.fr/climate/) averaged within the natural distribution range of *Quercus petraea* (red) and *Abies alba* (blue). Shaded area represents the standard deviation around the long-term average climate (1901–1960), prior to the acceleration of climate warming during the decades following 1960. The shapes of species distribution maps used to compute the temporal variation of the thermal niches were sourced from Euforgen ([http://www.euforgen.org](http://www.euforgen.org/)).

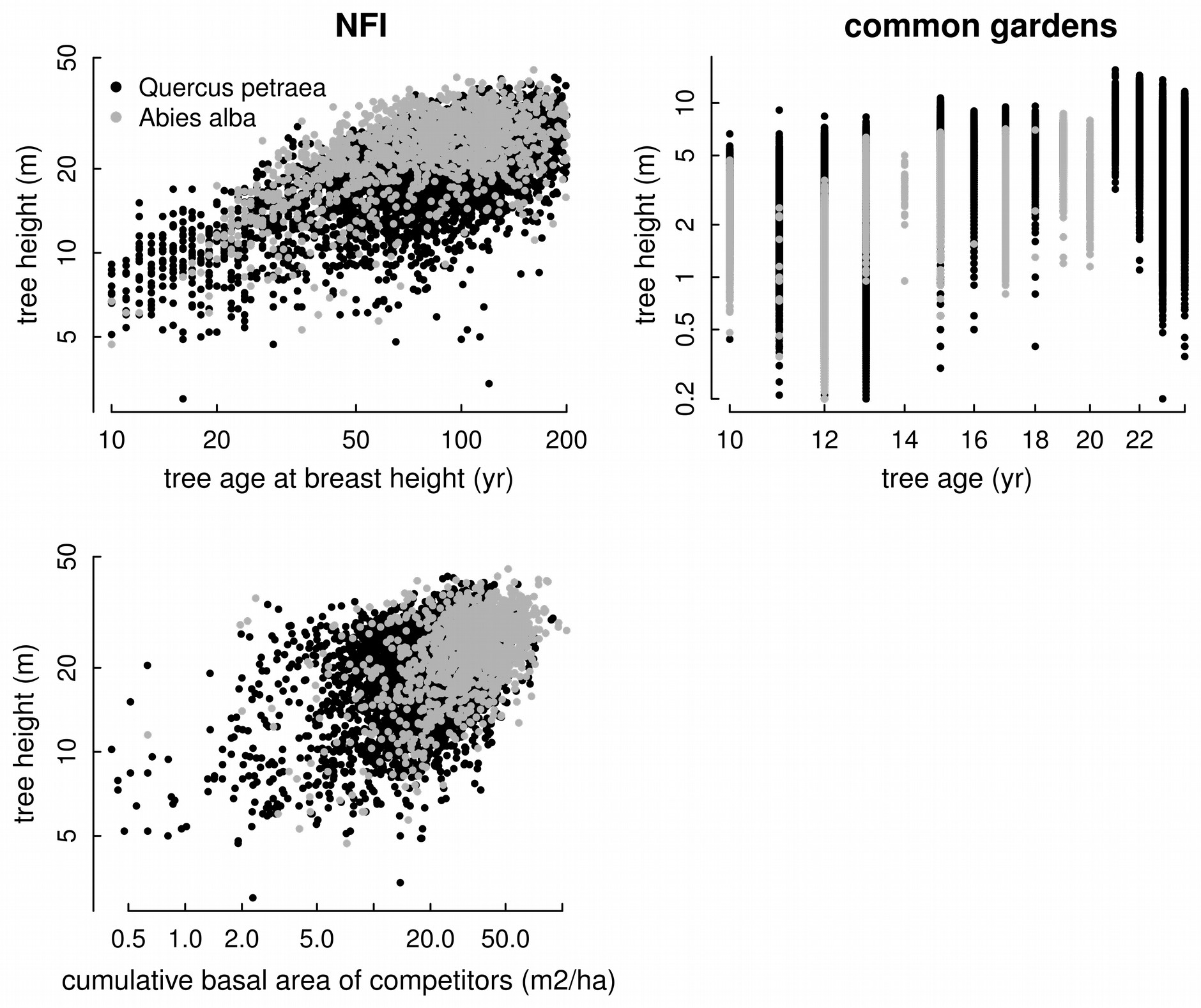

**Figure S4.1** Tree height variation as a function of tree age (top) and neighbour basal area (bottom) in *Quercus petraea* (black) and *Abies alba* (grey) in National Forest Inventories (NFI, left) and common gardens (right). Neighbour basal area was assumed to be constant between and within common gardens. Tree age was estimated using wood increment cores collected at breast height (1.30 m) in NFI plots, whereas tree age is the time between the height measurement and the sowing date in common gardens. Note a log scale on both axes.

**Table S1** Description of the 6 French common gardens used for measuring phenotypic traits on *Abies alba* provenances. Data include: test site code (as in Table S3), name of forest where the site is located, name of region where the site is located, latitude (in degree decimal, to the 4^th^ decimal point), longitude (in degree decimal to the 4^th^ decimal point), elevation (m a.s.l.), total size of the test site (in hectare), plantation density (trees per hectare) and number of *A. alba* provenances tested in the study (N).

| **Site code** | **Site name** | **Latitude** | **Longitude** | **Elevation** | **Planting date** | **Size** | **Density** | **N** |
| --- | --- | --- | --- | --- | --- | --- | --- | --- |
| 60301 | Bois Génard | 48.1167 | 4.9833 | 320 | 1972 | 3 | 2500 | 2 |
| 70201 | Rouvre Sur Aube | 47.0167 | 4.9667 | 410 | 1967 | 3.88 | 2500 | 21 |
| 70203 | La Brugère | 44.5667 | 3.4517 | 1110 | 1967 | 3.22 | 2000 | 16 |
| 70402 | Les Chauvets | 44.5667 | 3.4483 | 1050 | 1972 | 4.31 | 2500 | 20 |
| 70502 | Somail Chinchidou | 43.5333 | 2.7333 | 920 | 1973 | 0.29 | 10000 | 35 |
| 70503 | Somail Sagnassol | 43.5361 | 2.7347 | 973 | 1973 | 0.42 | 2500 | 33 |

Note: the test site may contain other genetic material than the populations tested in this study, e.g. other species irrelevant to this study.

**Table S2** Description of *Abies alba* Mill. provenances in the 6 common gardens. Data include: population code number and name, geographic coordinates in degree decimal, country of origin, and number of blocks (replicates) per site where the population is planted. ‘B&H’ Bosnia & Herzegovina, ‘Czech’ Czech Republic.

| **Pop. code** | **Pop. name** | **Latitude** | **Longitude** | **Country** | **60301** | **70201** | **70203** | **70402** | **70502** | **70503** |
| --- | --- | --- | --- | --- | --- | --- | --- | --- | --- | --- |
| 36549 | FORE | 45.517 | 3.550 | France |  | 5 | 5 |  |  |  |
| 36550 | GRBO | 45.350 | 4.517 | France |  | 5 | 5 |  |  |  |
| 36555 | RUHP | 47.767 | 12.650 | Germany |  |  |  |  | 80 | 32 |
| 36556 | BEAR | 42.967 | 2.400 | France |  | 5 | 5 |  |  |  |
| 36557 | RIAL | 42.950 | 2.367 | France |  | 5 |  |  |  |  |
| 36558 | NEBS | 42.900 | 2.133 | France |  | 5 | 5 |  |  |  |
| 36559 | PUIV | 42.900 | 2.033 | France |  | 6 | 5 |  |  |  |
| 36560 | CALL II | 42.850 | 2.117 | France |  | 5 | 5 |  |  |  |
| 36561 | FANG II | 42.833 | 2.283 | France |  | 5 |  |  |  |  |
| 36562 | LAFA | 42.767 | 2.000 | France |  | 6 | 5 |  |  |  |
| 36563 | BALC | 42.583 | 2.100 | France |  | 7 | 5 |  |  |  |
| 36564 | CANG | 42.550 | 2.450 | France |  | 6 | 5 |  |  |  |
| 36565 | JOUX II | 46.850 | 6.050 | France |  | 5 | 5 |  |  |  |
| 36566 | DONO | 48.517 | 7.167 | France |  | 5 | 5 |  |  |  |
| 36567 | BOAJ | 46.283 | 4.467 | France |  | 5 | 5 |  |  |  |
| 36568 | MOLL | 46.000 | 4.433 | France |  | 5 |  |  |  |  |
| 36569 | BONO II | 45.917 | 3.800 | France |  | 5 | 5 |  |  |  |
| 36570 | JASO | 45.533 | 2.133 | France |  | 5 |  |  |  |  |
| 36571 | ECOU II | 48.517 | 0.067 | France |  | 7 | 5 |  |  |  |
| 36572 | PERS | 48.300 | 0.300 | France |  | 5 |  |  |  |  |
| 36573 | LASU | 47.667 | 23.583 | Romania |  | 6 | 5 |  |  |  |
| 36574 | PRAH | 45.350 | 25.550 | Romania |  | 8 | 5 |  |  |  |
| 36575 | BLIZ | 51.117 | 20.750 | Poland |  |  |  |  | 80 | 32 |
| 36576 | LA-SU II | 45.850 | 22.483 | Romania | 5 |  |  |  | 80 | 32 |
| 36577 | KURN | 49.850 | 10.033 | Germany |  |  |  |  | 80 | 32 |
| 36578 | PRAZ | 44.483 | 7.050 | Italy |  |  |  |  | 80 | 32 |
| 36579 | KOZA | 45.000 | 17.063 | B&H |  |  |  |  | 80 | 32 |
| 36580 | FANG IV | 42.817 | 2.267 | France |  |  |  | 5 | 80 | 32 |
| 36581 | ZWIE | 49.017 | 13.233 | Germany |  |  |  |  | 80 | 32 |
| 36582 | CALL IV | 42.869 | 2.087 | France |  |  |  |  | 80 | 32 |
| 36583 | CAMA | 43.800 | 11.817 | Italy |  |  |  |  | 80 | 32 |
| 36585 | TRIE | 47.486 | 14.486 | Austria |  |  |  |  | 80 | 32 |
| 36586 | RYTR | 49.490 | 20.668 | Poland |  |  |  |  | 80 | 32 |
| 36588 | KOBE | 48.067 | 13.233 | Austria |  |  |  |  | 80 | 32 |
| 36589 | VODC | 44.135 | 17.189 | B&H |  |  |  |  | 80 | 32 |
| 36590 | KELH | 48.917 | 11.868 | Germany |  |  |  |  | 80 | 32 |
| 36591 | SEVR II | 48.791 | 0.464 | France |  |  |  | 6 | 80 |  |
| 36592 | JOUX III | 46.376 | 5.972 | France | 5 |  |  | 5 | 80 | 32 |
| 36594 | ROSE | 41.900 | 14.350 | Italy |  |  |  |  | 80 | 32 |
| 36596 | HOHE | 47.834 | 16.048 | Austria |  |  |  |  | 80 | 32 |
| 36597 | STSA | 49.563 | 20.636 | Poland |  |  |  |  | 80 | 32 |
| 36598 | SBRU | 38.583 | 16.333 | Italy |  |  |  |  | 80 | 32 |
| 36602 | BANS | 48.733 | 19.149 | Slovakia |  |  |  |  | 80 | 32 |
| 36603 | SLLU | 48.767 | 19.275 | Slovakia |  |  |  |  | 80 | 32 |
| 36604 | PODS | 49.283 | 20.183 | Slovakia |  |  |  |  | 80 | 32 |
| 36606 | FRYD | 50.921 | 15.079 | Czech |  |  |  |  | 80 | 32 |
| 36607 | LOCH | 47.457 | 7.640 | Switzerland |  |  |  |  | 80 | 32 |

**Appendix S2 Bioclimatic regionalisation of species natural distribution ranges**

In contrast to the common garden experiments, the geographical origin of the seed source (i.e., the provenance) is unknown for NFI observations. To overcome this major limitation, we made the neutral assumption that the trees have a local origin. In other words, we assume that potential seed sources were mostly from local provenances within the bioclimatic region with a low probability of long-distance transfer. We therefore performed a fine bioclimatic partitioning of the species’ natural distribution range (Figure S5.2). The long-term climate of origin (LTC) was estimated as the long-term (1901-1960) average climate of the bioclimatic region in NFI. The recent climate change (RCC) is estimated as the difference between the recent short-term climate (STC) experienced by the tree in the NFI plot (averaged over the last decade of growth before tree height measurement) and the estimated LTC for the corresponding climatic variable.

For each species, the natural distribution range (http://www.euforgen.org) was partitioned into bioclimatic regions by K-means clustering using the R package vegan (Oksanen et al. 2013) (Figure S5.2). We used the Caliński–Harabasz criterion (Caliński & Harabasz 1974) to select a statistically optimal number of regions (K = 70 for A. alba; K = 90 for Q. petraea) from a range of K (10 to 200 in steps of 10). Partitioning of bioclimatic regions was performed on Z-scores (standardised data) of long-term (1901–1960) averages of the following four climatic parameters: minimum temperature of the coldest month (°C), maximum temperature of the warmest month (°C), and total precipitation in the driest and the wettest months (mm); in addition to the latitude and longitude of the plots.

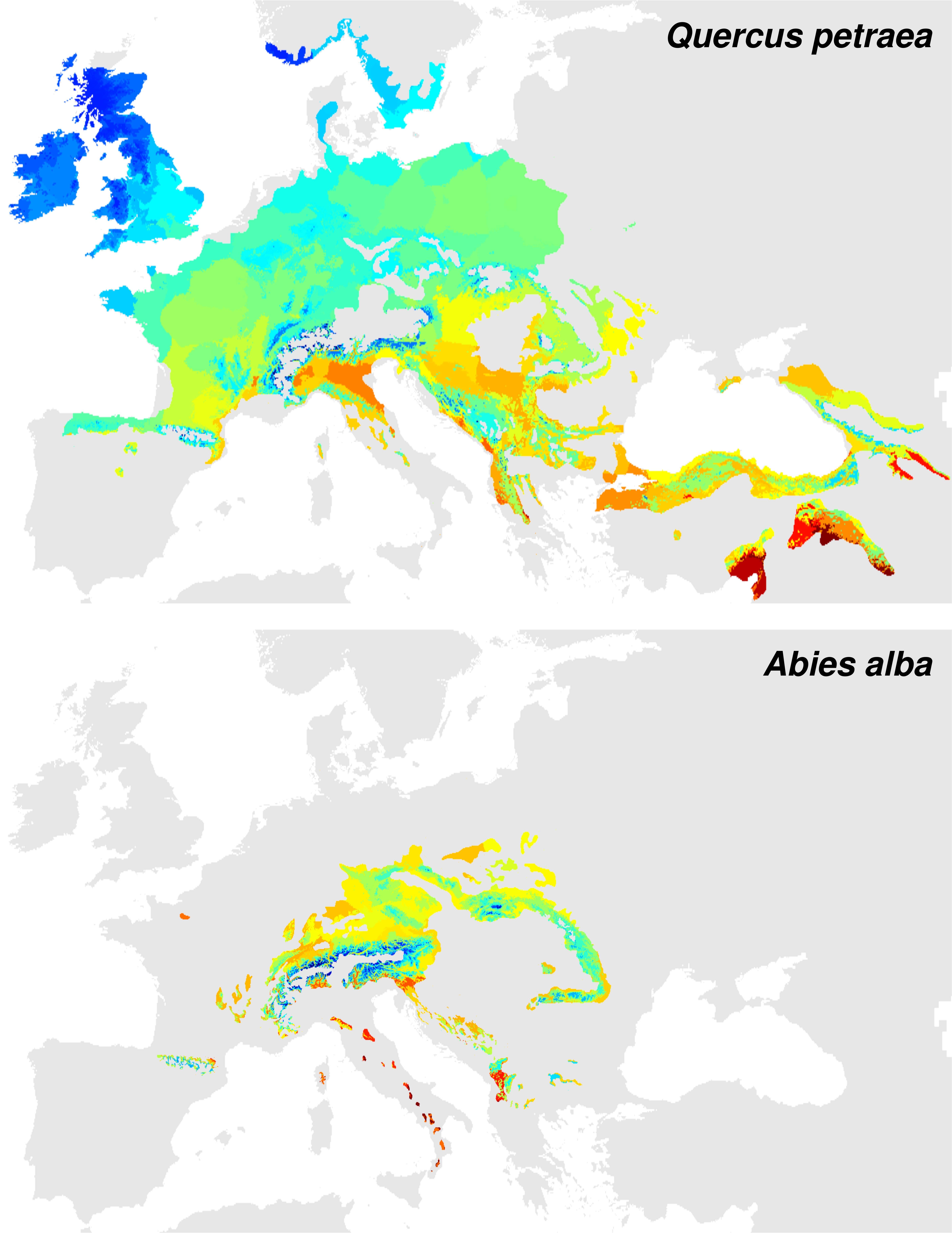

**Figure S5.2** Bioclimatic regions within the natural distribution range of *Quercus petraea* (top) and *Abies alba* (bottom). Colour gradient represents differences in long-term regional average (1901–1960) of annual mean temperature, from blue (coldest regions) to red (warmest regions). Discontinuous areas of similar colour indicate a similar long-term mean temperature between bioclimatic regions. Number of bioclimatic regions across species ranges: *N* = 90 and *N* = 70 for *Q. petraea* and for *A. alba* respectively.

**Appendix S3 Model validation**

*Validation using common garden data*

In the case of validation by comparing the predictions of *in-situ* models with raw common garden data, we standardized both predictions and raw data for differences in provenances, sites, age and neighbour basal area. First, the *in-situ* model was used to predict the mean height of each provenance in each site (‘provenance-by-site’ means) from equation 3 (main text), using the LTC of the provenance and the STC of the site (i.e. common garden) for a given age (12-year-old trees, i.e. the most common age at the time of height measurements across common gardens in both species) and neighbour basal area (30 m^2^ ha^-1^). Second, common garden data were standardized across ages using a linear mixed-effect model, where the log of tree height was regressed against the log of age with the provenance and the site set as random effects. In particular, we used model residuals computed from fixed effects only (i.e. age) to compute provenance-by-site means for common garden observations. Third, provenance-by-site means of common garden observations and *in-situ* model predictions were standardized as follows, before comparing them using Pearson correlation coefficients.

To estimate differences in tree height among provenances (provenance effect), provenance-by-site means were standardized (height values scaled between 0-1) independently for each site. In this way, we focused on height differences among provenances after accounting for differences in environmental conditions among sites (i.e. phenotypic plasticity effect). Then, we computed mean values by provenance. Pearson coefficients indicated significant correlations between common garden data and *in-situ* model predictions for the provenance effect in *Quercus petraea* (Fig. S6.3a) and *Abies alba* (Fig. S7.3a). Moreover, *in-situ* model predictions well predicted tree height differences among cold, core and warm provenances, as indicated by boxplots (Figs S6.3a and S7.3a).

To estimate plasticity, provenance-by-site means were standardized (height values scaled between 0-1) independently for each provenance. In this way, we focused on height differences among sites after accounting for the provenance effect. Pearson coefficients indicated significant correlations between common garden data and *in-situ* model predictions for the plastic component in *Q. petraea* (Fig. S6.3b) and *A. alba* (Fig. S7.3b). Considering cold, core and warm provenances separately, correlations were still significant at *P* < 0.05.

To estimate the total component of variation (sum of provenance effect and provenance plasticity effect), provenance-by-site means were standardized across all data (height values scaled between 0-1). Pearson coefficients indicated weak correlations in both species, significant in *Q. petraea* at *P* < 0.001 (Fig. S6.3c) and in *A. alba* at *P* < 0.10 (Fig. S7.3c). *In-situ* model predictions reasonably predicted height differences among cold, core and warm provenances, as indicated by boxplots (Figs S6.3c and S7.3c).

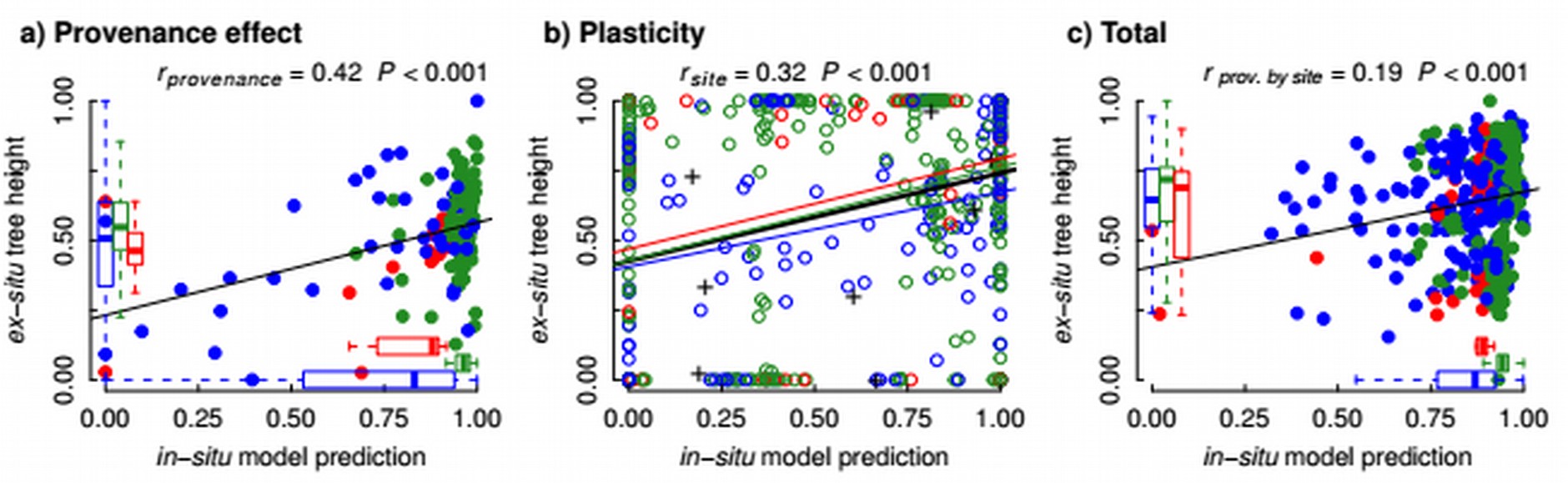

**Fig S6.3** Pearson correlation coefficients between *in-situ* model predictions (calibrated on NFI) and common garden data estimates of provenance (a) and plasticity effects (b) and of the total component of tree height variation (c) across *Quercus petraea* provenances. Points represent provenance means in a) and provenance-by-site means in b) and c). Tree height differences among cold (blue), core (green) and warm provenances (red) for common garden data and *in-situ* model predictions are indicated by boxplots in a) and c). Regression lines in b) illustrate correlations for phenotypic plasticity between common garden data and *in-situ* model predictions for cold (blue), core (green), warm (red) and all provenances (black); all are significant at *P* < 0.05; crosses indicate mean site values.

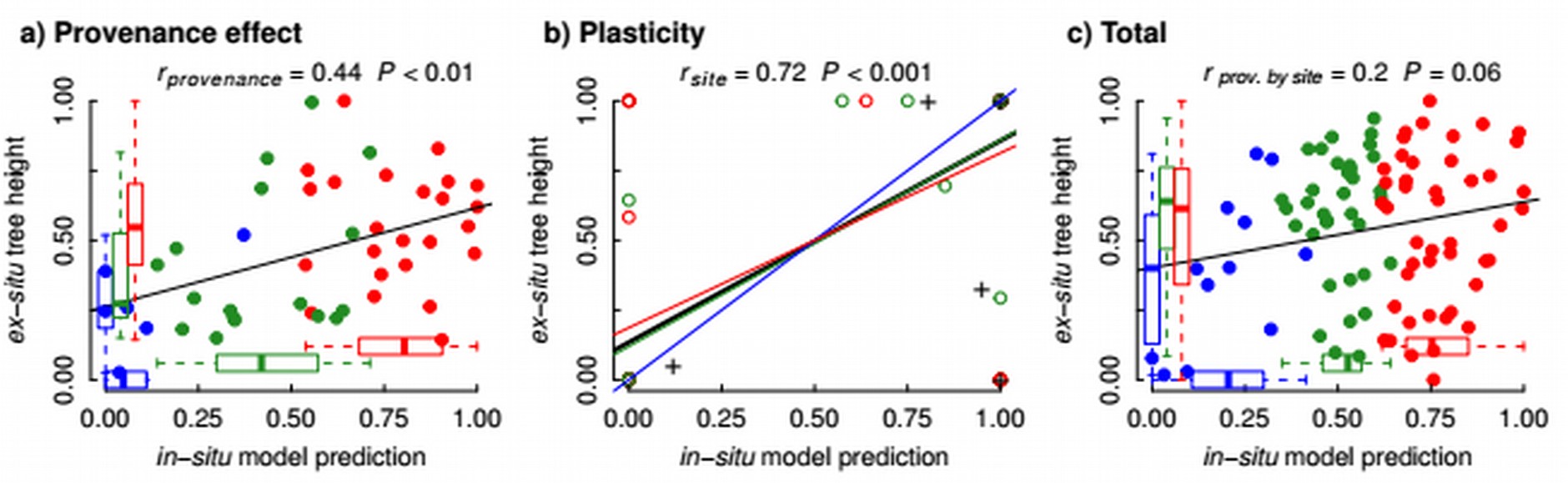

**Fig S7.3** Pearson correlation coefficients between *in-situ* model predictions (calibrated on NFI) and common garden data estimates of provenance (a) and plasticity effects (b) and of the total component of tree height variation (c) across *Abies alba* provenances. Points represent provenance means in a) and provenance-by-site means in b) and c). Tree height differences among cold (blue), core (green) and warm provenances (red) for common garden data and *in-situ* model predictions are indicated by boxplots in a) and c). Regression lines in b) illustrate correlations for phenotypic plasticity between common garden data and *in-situ* model predictions for cold (blue), core (green), warm (red) and all provenances (black); all are significant at *P* < 0.05; crosses indicate mean site values.

*Validation using ex-situ model predictions*

In the case of validation by comparing the predictions of both models, we predicted provenance and plasticity effects, and their interaction, as a function of LTC and STC conditions in common gardens. In particular, using equation 3 (main text) we predicted the relative variation of tree height as a function of *LTC_j_* (i.e., the provenance effect) by fixing *STC_k_* to mean observed values across common gardens (i.e., for average climate of planting sites). Reciprocally, the relative variation of tree height as a function of *STC_k_*  (i.e., the phenotypic plasticity effect) was fitted by fixing *LTC_j_* to mean observed values across provenances (i.e., for a mean-climate provenance). To predict plastic responses of each provenance *j*, we fitted height as a function of *STC_k_* and the interaction term *LTC_j_* × *STC_k_*. For this, we scaled predicted values between 0–1 independently for each provenance *j* to focus on the relative variation of height among provenances. The total component of variation was fitted as a function of *LTC_j_*, *STC_k_* and *LTC_j_* × *STC_k_*, i.e. the sum of provenance and plasticity effects and their interaction. Covariates in equation 2 (main text) were fixed to constant values, i.e. *age* (the most common age in common garden data, 12-year-old trees), *BAc* (mean observed value in NFI, ~30 m^2^ ha^−1^) and their interaction with *RCC_jk_* and *LTC_j_*, respectively. Confidence intervals (SD) of predicted values along *STC_k_* and *LTC_j_* gradients were computed by bootstrapping, i.e., 200 model runs on 50% randomly sampled trees with replacement. In the comparison of plasticity among provenances, confidence intervals (SD) of predicted values along *STC_k_* were computed among ‘cold’, ‘core’ and ‘warm’ provenances.
